# Four-dimensional omics data reveals ribosome heterogeneity, regulation of translation efficacy, and nonsense-mediated decay in the differentiation of spermatocyte to round spermatid

**DOI:** 10.1101/2023.07.13.548784

**Authors:** Szu-Shuo Lee, Ying-Chih Kung, Yuh-Shan Jou

## Abstract

A protein expression is regulated by transcription, translation, and sequential processing. However, well correlated RNA and protein abundance just only proportionate 40%, and even poorer when cell was stressed, differentiated, or tumorigenic transformed. Here, we discovered spermatocyte (SP) differentiated to round spermatid (RS) had equal regulation extent which may related to ribosomal behavior alteration. The change of ribosome occupancy was related to SP and RS specific function in spermatogenesis. Interactome of functional ribosome in SP and RS revealed the activated ribosome in SP but stalled and nonsense-mediated decay (NMD) associated ribosome in RS. Functional ribosomes of RS occupied 5’untranslated regions (5’UTR) of SP specific transcripts and correlated its’ RNA and protein downregulation. These findings suggested a branched NMD pathway was activated in RS to eliminate SP specific transcripts and keep them from being translated. Our discovery suggested the heterogeneity of ribosomal interactome may play an important role in spermatogenesis.

## INTRODUCTION

In multicellular organisms, knowing the molecular basis of phenotypes under cell fate commitment at systems-level is a fundamental and critical question to understand how the spatiotemporal gene expression to orchestrate the dynamic molecular and morphological changes for keeping biological processes on time during development[1]. For a specialized cell, subsets of necessary genes should be expressed at the proper time and subcellular localization at both RNA and protein levels to conduct the assigned physiological functions[2]. Therefore, expression of necessary genes, but not global expression, is regulated at multiple levels such as epigenetic, transcriptional, post-transcriptional, translational and post-translational modulations in response to genetic and environmental stimuli[3]. Although majority of gene expression is controlled at transcriptional level, pervasive transcription of genomic DNA further suggested that tightly regulatory mechanisms at the post-transcriptional, translational and post-translational levels are required to monitor the quality of gene expression for removing unnecessary and aberrant transcripts and polypeptides to ensure expressing proper proteins at the appropriate times and levels[4]. With recent advances of various deep-sequencing and proteomic technologies accompanied with integrated omics analysis, exploration of gene expression with integrated analysis of transcriptome, translatome (ribosome foot printing) and proteome (quantitative mass spectrometry) results at systems-level across different committed cell types is lagging behind for better understanding the cellular determinants for driving the proper biological function in specialized cell types[5-8].

Spermatogenesis is a continuous cell differentiation process from spermatogonia developing to spermatozoa (or sperm cells) in the seminiferous tubules of the testis. The entire process is tightly regulated for sperm maturation from diploid spermatogonium divides mitotically to two primary spermatocytes [9]. The spermatocytes will duplicate genome and undergo meiosis I to produce two diploid secondary spermatocytes followed by meiosis II to divide into four haploid spermatids. The spermatid will go through an elongated transformation process called spermiogenesis to allow the growth of microtubules bundled into axoneme and form a tail for sperm motility. During spermatogenesis, widespread transcription of genomic DNA was transcribed and RNA granules such as the inter-mitochondrial cement (IMC) and the chromatoid body (CB) were observed starting in spermatocytes [10]. Subsequently, elongated spermatid genomic DNA is packaged and gradually replaced by nuclear basic protein protamine to form the highly condensed and transcription inactive chromatin featured with translational repression. As aforementioned, with increasing evidence to show the pivotal roles of post-transcriptional and translation regulation of gene expression in the specialized germ cells, we selected murine spermatogenesis as a model and applied integrated approaches of transcriptome (i.e. RNA-seq), translatome (i.e. Ribo-seq) and proteome for a systems-level of understanding the specific genes expression from meiotic spermatocytes to post-meiotic haploid round spermatids for conducting their designated functions.

With recent advanced omics technologies, post transcriptional and translation regulation is generally believed to become the dominant sources of modulating gene expression to fulfill proper biological functions and response to altered environmental stimuli in specialized cell types after comparisons with naïve parental cell types. With availability of transcriptome and translatome datasets in the public domain for mouse spermatocytes and round spermatids, we performed integrated omics analysis and ribosome interactomes to illustrate the heterogeneous profiles of ribosome associated proteins guided transcript-specific translation in mouse spermatocytes and spermatids. Interestingly, we identified different nonsense-mediated decay (NMD) factors associated with functional ribosome interactomes in spermatocyte and round spermatid. Together, after validations with cell-type specific gene expression selected from spermatocyte and round spermatid, our results suggested detail molecular basis of post-transcriptional and ribosomal modulation of cell type specific gene expression in systems-level reflecting mechanisms of cell fate determination in spermatogenesis.

## RESULTS and DISCUSSION

### Comparison of transcriptional and translational concordance of gene expression in specified cell types undergoing differentiation and oncogenesis

To gain a systems-level view of gene expression modulation in between transcription and translation during multiple biological processes, since there might have other biological processes with transcriptional-translational conflict which same as in tumorigenic suppression of bladder cancer[11]. we collected transcriptome (TR, i.e. RNA-seq) and translatome (i.e. Ribo-seq) datasets of the same cell lineages (Supplementary Table1) and investigated large scale transcriptional-translational conflict or concordance in regulation of gene expression under stimuli of differentiation and oncogenesis (Fig. 1) [12-18]. In general, the comparisons of gene expression modulation dominant either at transcription and/or translation stage during transition of cell fate changes are based on pairwise comparison of histological primary/parental/ground state, intermediate state, and differentiated/activated/transformation state of specified cell types. For instance, we compared different stage cells in spermatogenesis and embryonic stem cells versus matured cell types were considered as developmental differentiation. On the other hand, Kras mutant driven quiescent, pre-senescent, senescent, tumorigenic transformed fibroblast, and overexpressed MYC in cell lines were categorized as oncogenic stimuli.

**Fig 1.**
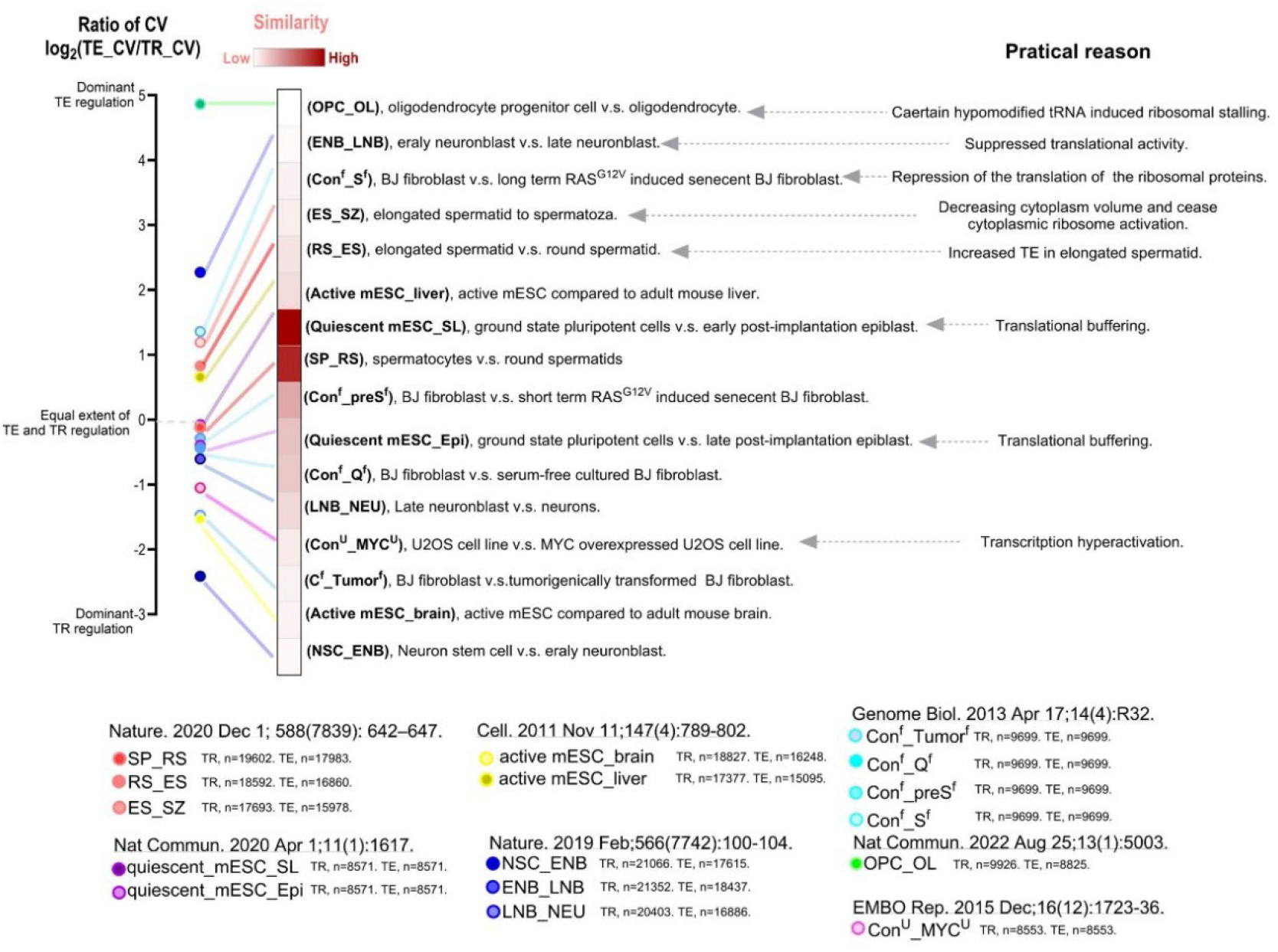
Extent of regulation between translation and transcription in the spermatocyte/round spermatid is similar pluripotency cell status. To narrative range of transcription regulation and translation regulation, coefficient of varication (CV) was used to compare extent of these two regulations. CV ratios were counted by dividing CV of TE to CV of TR. 6 dataset were used to plot and OPC_OL as the one has highest extent of translation regulation. In the other hand, NSC differentiate into neuroblast showed highest extent of transcription regulation. Differentiation of quiescent mESC into cell lineage were known related to ribosome buffering of translation. Similarity indexes were calculated from converting log 2 based "Ratio of CV” to absolute value and inversing it into reciprocal for further indication of the proportionate extent from TR and TE. Practical reasons were extracted from the conclusion of the given papers. CV: Coefficient of variation; TR: Transcription; TE: Translation efficiency; Translation efficiency (TE)/Ribosome occupancy in CDS regions (R.O. in CDS) = Ribo-seq reads in CDS region/ RNA-seq reads in CDS region (from cytosol); and Similarity index: 1/|log2(Ratio of CV) |. SP: spermatocyte; RS: round spermatid; ES: elongated spermatocyte; SZ: spermatozoa; OPC: oligodendrocyte progenitor cell; OL: oligodendrocyte; NSC: neural stem cell; ENB: early neuroblast; LNB: late neuroblast; NEU: neuron; active mESC: mouse embryonic stem cell without LIF medium; quiescent mESC: mouse embryonic stem cell with serum free medium, GSK inhibitor and MEK inhibitor;SL: mouse embryonic stem cell with serum/LIF medium, activated relatively. Epi: Epiblast stem cell, differentiated from ESC; C: BJ fibroblast Tumor: BJ fibroblast which transfected with oncogenic KRAS, p16INK4A knockdowns and SV40 small-T expression and acquired tumorigenic function; Q: BJ fibroblast with quiescence induced by serum depletion; preS: senescence induced BJ fibroblast which transfected with oncogenic KRAS in short period. S: senescence induced BJ fibroblast which transfected with oncogenic KRAS in long period; Con: Parental U2OS cells; Myc: U2OS with MYC transfection.

To rank the dominancy of gene expression modulation from transcription or translation, the coefficient of variation (CV) was applied to narrative the modulation intensity. By comparing the CV of translation efficiency (TE_CV) versus CV of transcription (TR_CV) of 16 pairwise comparisons of cell fate changes in either differentiation or oncogenesis. Consistent with previous reports, our results demonstrated that oligodendrocyte progenitor cell (OPC) differentiation to oligodendrocyte (OL) experienced ribosome stalling due to shortage of at least 2 tRNAs in oligodendrocyte. In contrast, neural stem cells (NSC) differentiated into early neuroblasts (ENB) had a higher extent of transcription regulation than that of translation regulation. Since higher TR_CV than TE_CV is a proxy measurement for reflecting gene expression status in primary cell fate commitment: mESC compared to brain which is a terminal differentiated neuron organ, NSC differentiated to ENB, or oncogenic processes: MYC overexpressed cells and tumorigenic transformation. Inversely, higher TE_CV than TR_CV was more related to cell fate decided status or cellular status: OL differentiation, spermiogenesis, and senescence. Further, our results indicated that differentiation process from spermatocyte to round spermatids (SP_RS) had almost equal ratio of transcription and translation regulation with comparison to that in the ground state to primed pluripotency (Quiescent mESC_SL) statuses (-0.11698 and -0.07731, log 2 based value). Translational buffering, ribosome translating is shifted from a subset of transcripts to another subset of transcript without change proteins expression, was indicated in the differentiation from zygote to post-implantation. This made us want to investigate whether SP_RS have shifting of ribosome translation as the life beginning after fertilization.

TE is calculated by ribosome occupied regions in CDS (coding DNA sequences) region and normalized with RNA abundance from RNAseq. Ribosome occupancy might not only suggest ribosome translating to the given transcripts but also could be related to slow or stop translating ribosomes. Due to the ambiguous of the definition the situation of translation, it had been judged had poor correlation to proteome or correlated well in subset of genes/transcripts only[19], it implicated that the complexity of post-transcriptional and post-translational regulation. Here, we change the wording from TE to R.O (stands for absolute ribosome occupancy, Riboseq reads normalized by RNAseq reads in coding DNA sequences) to specify the role of ribosome behavior.

### In SP-RS period, RNA transcription could positively correlate to biological functional proteome of RS, and R.O. in CDS negatively or non-significantly correlate to biological functional proteome of SP

To invest in what proteins were dominantly regulated by transcription or R.O. in CDS region, the majority of differential expressed proteins (DEPs) between SP and RS were categorized by Biological Process in Gene Ontology terms (GOBP) (Supplementary Table2 and supplementary Figure 1) and classified by difference of TR and R.O in CDS from SP and RS comparison statistically. The majority differential expressed protein categories by Gene Set Enrichment Analysis (GSEA) with GOBP were shown (Supplementary Figure 2A,2B). Each of the 4 dominant and well-known related to SP or RS categories were selected and classified into 9 groups. With comparing the protein expression with certain regulation pattern, we found that the protein expression correlated to “GOBP CELL CYCLE”, “GOBP CHROMOSOME ORGANIZATION”, and “GOBP DNA METABOLIC PROCESS” did not significantly correlated into specific groups, except “GOBP ORGANONITROGEN COMPOUND BIOSYNTHETIC PROCESS”. Proteins belonging to “GOBP ORGANONITROGEN COMPOUND BIOSYNTHETIC PROCESS” showed less R.O. in CDS in SP were correlated to higher protein expression than other groups (Figure 2). This might be the similar scenario which had been found in the ground state-to-primed pluripotency statuses, shifting of R.O. in CDS but not correlate to higher protein expression. In contrast, the proteins in all four categories belonged to RS were shown more transcription in RS was correlated to higher proteins. We concluded that when SP transited to RS may have undergone a ribosome-participant regulation which may selectively regulate subset of transcripts by R.O. in CDS alteration but not correlating to nascent peptides synthesis, such as cell cycle related ones.

**Fig 2.**
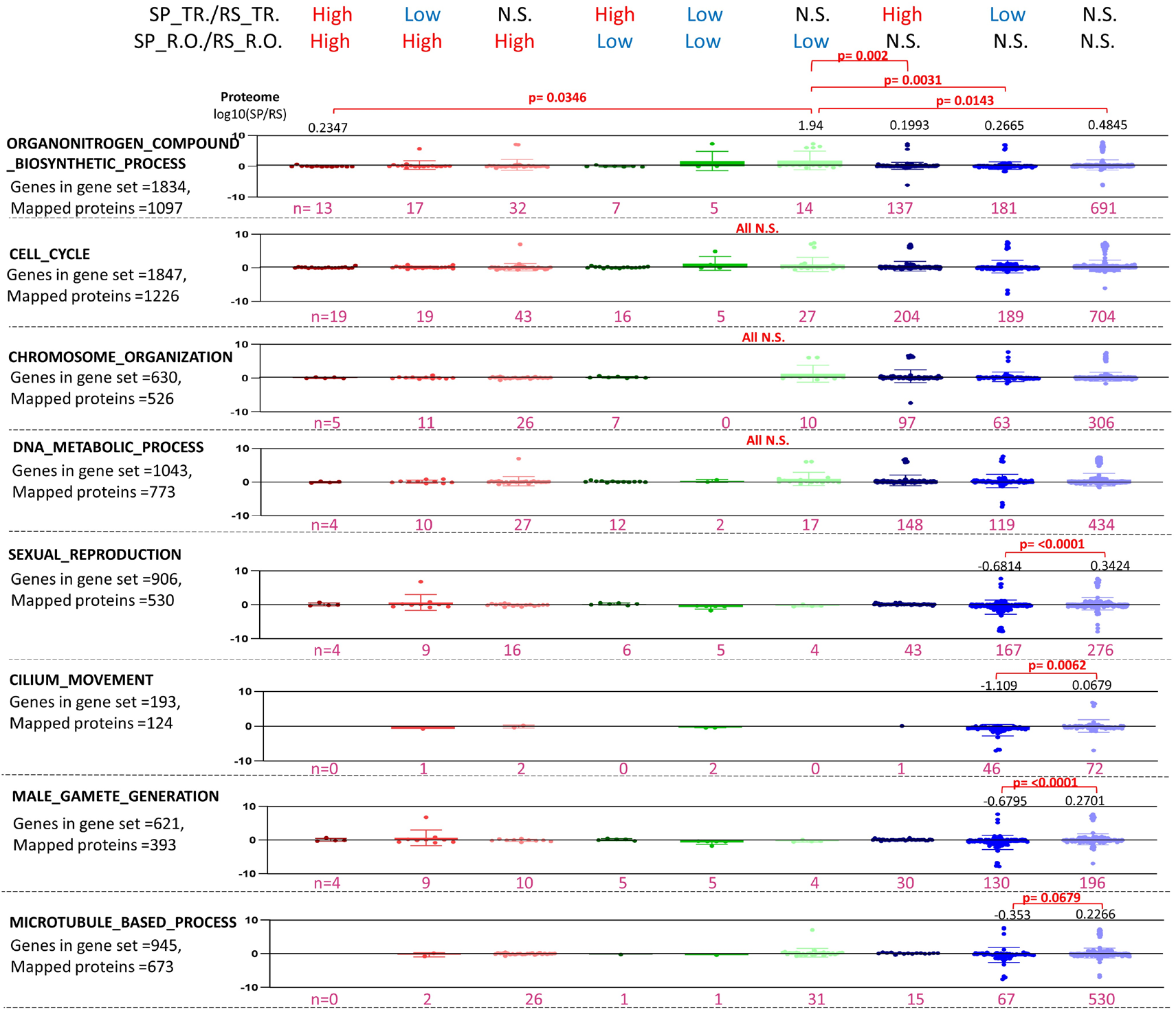
Relatively low R.O. in SP correlated to high protein expression, and relatively high transcription in RS correlated to high protein expression. RNA expression level change from SP to RS (SP_TR./RS_TR.) and R.O. level change from SP to RS (SP_R.O./RS_R.O.) were separated into 9 groups. Differential protein expressions (SP/RS value with log 10 based transformation) of certain protein from each of the 4 most significant SP and RS related biological processes were annotated by dot plots. The number with pink color demonstrated the gene number in each category. The mean value was annotated above the dot plot when the comparison was significant. One-way ANOVA and Turkey’s test was applied for statistical analysis for differentiation between each category.

### R.O. in CDS was non-correlated to RS proteome

To further address the correlation of proteome between transcriptome, ribosome foot printing, and R.O. in CDS with global view, the result had been shown by heatmap (Figure 3A). Transcriptome and ribosome foot printing were with significantly higher correlation with proteome instead of R.O. in CDS. This could be explained by ∼60% of genes are correlated to RNA expression or transcriptome[15, 20]. Ribosome foot printing retains the characteristics of transcriptome pattern and correlates to proteome better than R.O. in CDS value which has been normalized by RNA expression. R.O. in CDS could reflect the information of ribosome behavior more specifically than ribosome foot printing results. The RS cells had the poorest correlation between proteome and R.O. in CDS (Figure 3B). To investigate the regulation extent of TR and TE, all five spermatogenic cells were compared (GC1 cell line, which was immortalized by SV40 early region, was considered as spermatogonia). The result indicated more TE regulation was relied on along with spermatogenesis process (Supplementary Figure 2). It reflected the more condensed nucleus in spermatogenic cells undergoing more TE regulation. This suggests that RS cells, which was the last cellular stage right before the magnitude of nucleus condensing was enough to global suppressing transcription, may have different ribosome behaviors or regulation in this cellular status of spermatogenesis.

**Fig 3.**
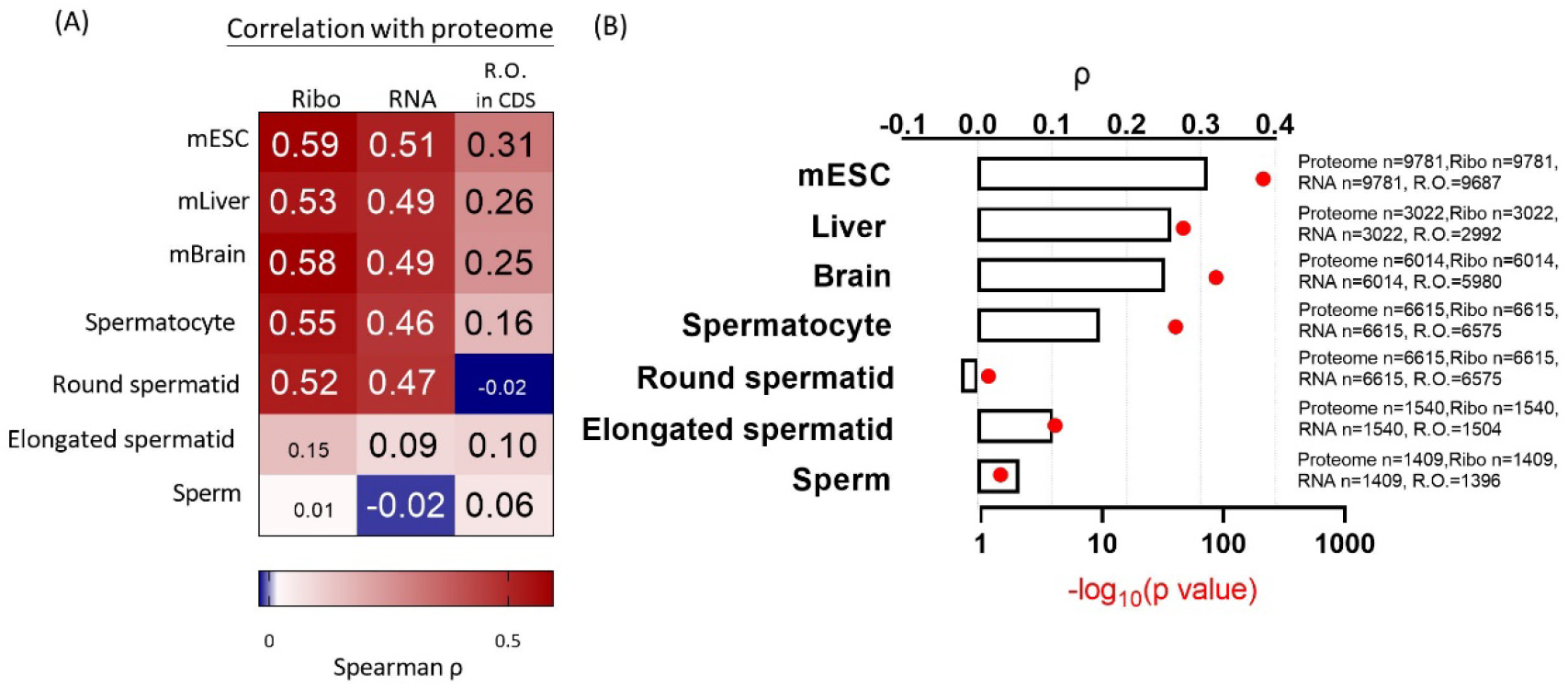
Translation efficiency is not well correlated to proteome in spermatogenesis process. (A)The translatome, transcriptome, and R.O. in CDS region were calculated and determined the correlation of the corresponding proteomes. Spearman rho value were showed in left heatmap, and statistical significance by font size. (B) Extract R.O. value in CDS and p-value to determine the differences between the cells. The gene number in each omic data was annotated on the left.

### RAPs of functional ribosomes determined SP had relatively translationally activated functional ribosome and RS had activated NMD factors binding functional ribosomes

As known that regulation of R.O. in CDS was altered between SP and RS, and ribosome interactome (Ribosomal associated proteins, RAPs) could cause R.O. pattern and protected fragments altered in different context [21]. We dissected the interactome of functional ribosome in isolated SP and RS from mice. By targeting Rpl36, which majorly be assembled into functional ribosomes[22], the functional ribosome interacted proteins were identified and quantitated. Pull-down protein quantity was normalized for further comparison (Supplementary Figure 3A, Figure 3B). Dissect the interactome analysis by GSEA showed ribosomal components and RNA binding proteins were the most two significant groups, which should be the most two large groups of ribosomal interactome[22], in the Rpl36 pull-down proteins (Figure 4A). For further investigate how different between functional ribosome interacted proteins between SP and RS, PCA analysis (Figure 4B) and differential protein abundance scatterplot (Figure 4C) indicated SP and RS have different pattern of functional ribosome interacted proteins, suggesting ribosome heterogeneity between SP and RS. Furthermore, functional ribosomes in SP interacted with more proteins than RS. This result might reflect previous findings that RS has poor protein synthesis and low TE[18, 23]. Enrichment of eIf2 families and positive regulators of translation proteins in SP functional ribosome interactome also support this finding (Figure 4D). With the aspect from SP to RS may went through ribosome occupancy shift like the ESC from ground state-to-primed pluripotency statuses, we addressed two significant differences of translational activator and RNA Nonsense-Mediated Decay (NMD) factors, Stau1 in SP functional ribosome interactome and Upf3b in RS functional ribosome interactome. Stau1 protein facilitates translation by stabilizing RNA and also the key factor to regulate RNA decay with non-classic NMD, Stau1-mediated mRNA decay (SMD). In the other hand, Upf3b is a major factor in NMD. Here, we suggested that alteration of translation activation and RNA decay may be respond to poor correlation between R.O. in CDS region and proteome when SP transited to RS in spermatogenesis process. Furthermore, because functional ribosomes were interacted NMD based factors Upf1, Upf2, and Upf3A both in SP and RS, the RNA decay mechanism between SP and RS may different, SMD and NMD for instance, and targeted different transcripts, like in myogenesis and certain cell types[24, 25].

**Fig 4.**
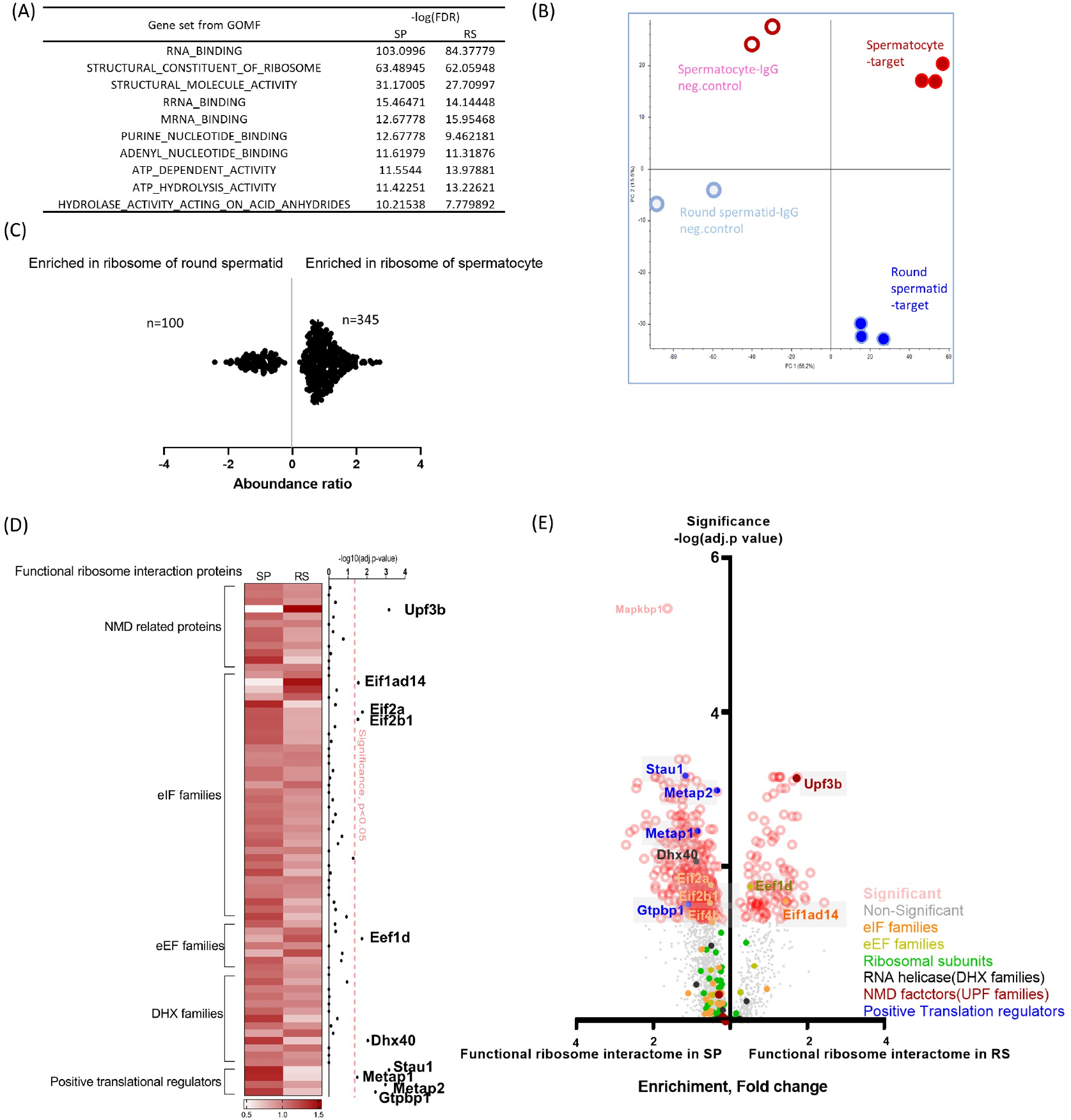
Ribosome in RS may be deficient in translation initiation and prime to RNA NMD process. (A) Narrative of interactome from SP and RS’s functional ribosomes by GSEA. (B) Functional ribosome associated proteins were revealed by Rpl36 immunoprecipitation of spermatocytes and round spermatids. The result was presented by PCA plot to determine the distinguish. (C)Functional ribosome associated proteins which significantly enriched in spermatocytes and round spermatids were indicated by abundance. (D) Revealing functional ribosome interactome in spermatocytes and round spermatids. The heatmap was categorized by the function of majority differential interacting proteins. A significant adjust p-value was plotted in the left of heatmap individually. (E) Volcano plot of total functional ribosome interacted proteins.

### R.O. in 5’UTR and 3’UTR negatively correlated to RS proteome

Since NMD factor Upf3b dominantly interacted with functional ribosomes in RS cells, moreover, functional ribosomes translate on certain transcripts would be the key step to activate NMD on this transcript[26]. We investigated the R.O. from different regions of transcripts and the correlation of proteome. The considerable significant negative correlation between proteomes and R.O. in 5’UTR and 3’UTR which from 5’UTR and 3’UTR and in SP and specifically in RS (Figure 5A-C). This result corresponds to the data from interactome of functional ribosome, NMD factors interacted with functional ribosome in RS cells. NMD could be the mechanism to eliminate transcripts from previous status, such as SP transit to RS, RS need to reshape the transcriptome and proteome and get rid of the rest of transcripts and proteins from SP. Published references have indicated ablation of the NMD factor and complex of E3 ubiquitin ligases would dampen developmental process such as neuron maturation and spermatogenesis[27-31]. Here, we addressed R.O. from different transcript regions in RS by enrichment analysis which classified by GOBP terms composed from SP and RS proteomes (Figure 5D). With calculation enrichment score from ssGSEA[32], we could calculate the enrichment of given R.O. value of cells in proteome GOBP categories. Enrichment score from value of SP R.O. or RS R.O. in 5’UTR, CDS, and 3’UTR represented a translation-like pattern (highest score in CDS compared to the scores from 5’UTR and 3’UTR) in its own proteome categories. Additionally, the enrichment score from SP R.O. in RS proteome categories showed that SP R.O. also correlated to most of categories with translation-like pattern. However, the Enrichment score from RS R.O. in SP proteome categories showed that RS R.O. correlated to most of categories but without translation-like pattern. (Supplementary Figure 4A-B, Figure 5D). This result supported the hypothesis we argued that the RS R.O. to some degree was related to the NMD activation. Because of NMD activation, the RS R.O. in CDS had no correlation to the proteome. 5’UTR and 3’UTR were significantly negatively correlated with the proteome. The SP related functional transcripts would be the majority of the RS ribosome mediated NMD targets. The NMD pathway in RS may not classical NMD pathway but induced by branched NMD pathways[33]. In other hand, the enrichment scores of SP R.O. had translation-like pattern in both SP and RS GOBP categories of functional proteins was corresponding to previous discovery that SP had transcribed and translated RS related proteins minorly[34, 35].

**Fig 5.**
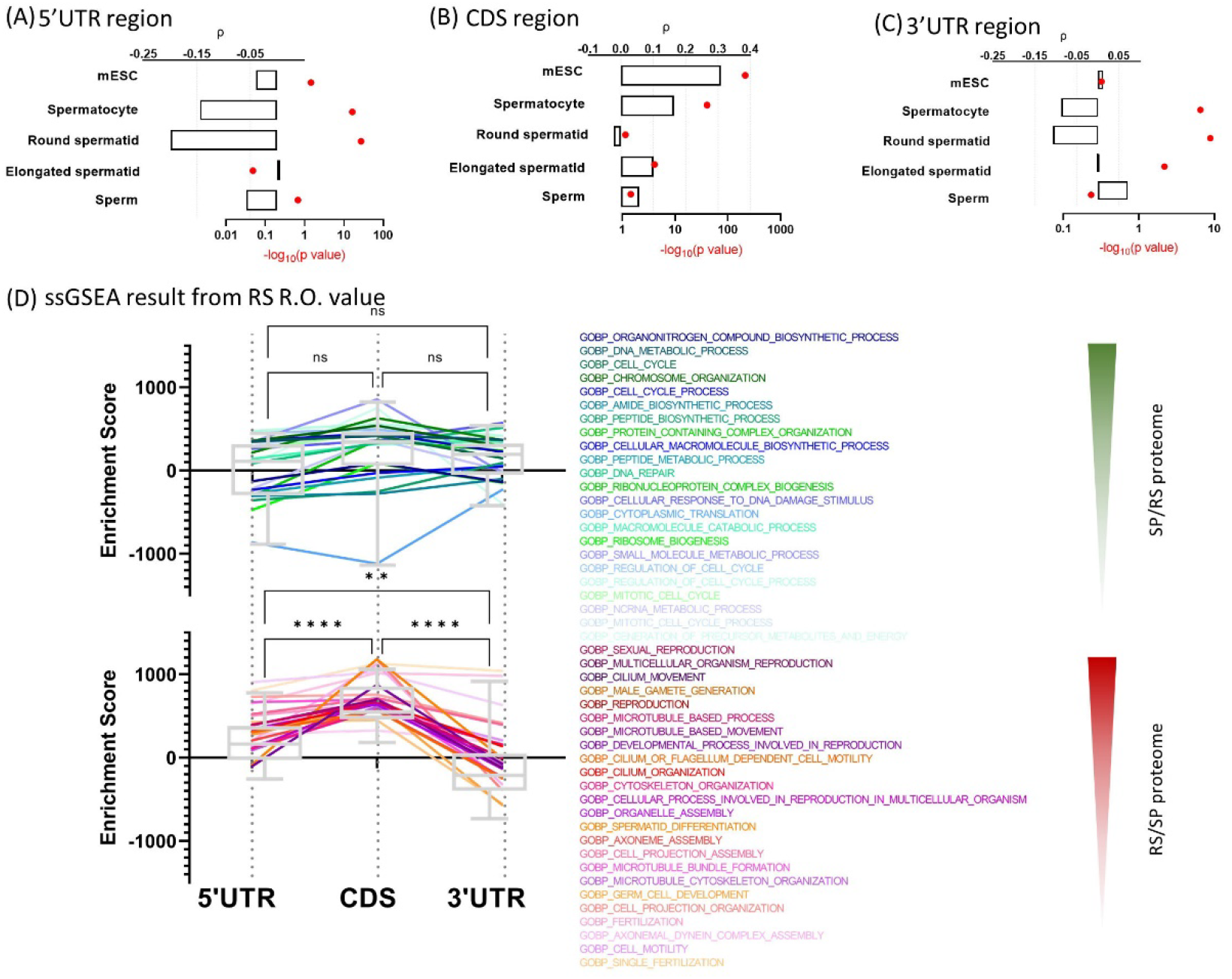
Ribosome occupancy (R.O) of 5’UTR and 3’UTR in RS status showed negative correlation to protein expression, which were functional proteins in SP. Ribosome occupancy (R.O) was calculated by translatome divides by transcriptome in the given regions. (A) 5’UTR regions, (B) CDS regions, and (C) 3’UTR regions were extracted specifically and calculated the correlation of corresponding proteome. (D) Enrichment score of ssGSEA revealed the correlation of R.O values in different region and the functional categories from SP or RS dominant proteome.

### RS had equal R.O in 5’UTR of SP specific transcripts as SP had

To be more specific, addressing the difference ribosome binding shifts between SP and RS transcripts to support our assumption, we compared the SP R.O and RS R.O. on different regions of SP specific transcripts and RS specific transcripts. As most ribosomes were located at CDS, SP R.O. and RS R.O. were both relatively higher than the other. However, as previous RS R.O. in 5’UTR negatively correlated to its proteome, RS R.O. in 5’UTR were equal as SP R.O. in 5’UTR in SP specific transcripts and even higher than SP in RS specific transcripts (Figure 6A-B). This suggested that ribosomes in RS may have had different behavior compared to it in SP. The results we argued here that the reinforced branched NMD feature of functional ribosome and dampened ribosomal function of regular translation in RS, and the scenario of 5’UTR seemed may have the distinguished role in this novel mechanism in the differentiation of SP to RS.

**Fig 6.**
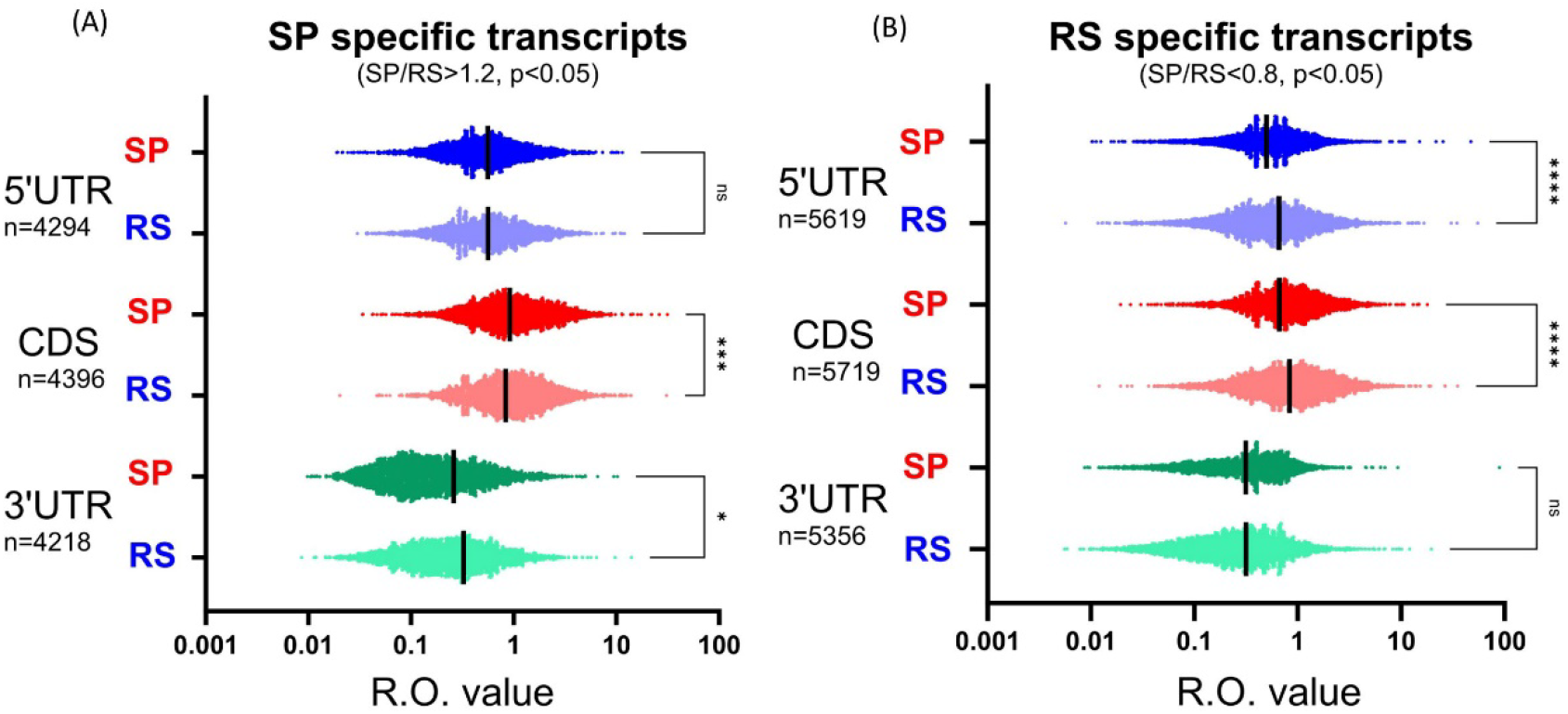
RS R.O. in 5’UTR of SP specific transcripts were similar to SP. SP and RS specific transcripts were distinguished by fold change and significance. R.O. values in different regions from SP and RS were calculated and annotated by dot plots. (A)Result in SP specific transcripts, and (B) result in RS specific transcripts.

### Sycp3 and other SP specific proteins could be cleared in RS by NMD to corresponding transcripts through R.O in 5’UTR

Seeking more evidence for our investigation, SP and RS well known proteins picked to as demonstration. To validate the R.O. shifting between SP and RS in these critical genes/transcripts, Sycp3 [36, 37]and Top2a[38, 39] were participated in meiosis and selected as SP specific genes, Ldfc[40], Pgam2[41] and Tubb4b[42] were component of mature sperm and choose as RS specific genes. Clu protein[43] were choosing as internal control which expressed both in SP and RS. In Sycp3 and Top2A transcripts, which majorly translated in SP, the ribosome still binding at 5’UTR and CDS when transited to RS, although the R.O. were decreasing when comparing to SP. However, the binding pattern, RS R.O. in 5’UTR and CDS were almost equal, was not similar to the SP pattern which related to regular translation. In the other hand, RS specific transcripts have equal R.O. pattern between SP and RS. Due to Ldhc transcripts being transcribed in SP status, Ldhc may transcribed and translated simultaneously, and the quantitation of proteins were accumulated through SP to RS transition. Intriguingly, one of Sycp3 transcripts was annotated be classic as NMD substrate had significantly highest R.O. in 3’UTR, which is the exon-junction complex dependent canonical NMD (≥50-55nts downstream of premature stop codon[44]) happened locus. Our additional finding on 5’UTR gave us more confidence that R.O. increase in 5’UTR were related to eliminating transcripts of prior cell status by a branched NMD mechanism in SP to RS transition.

Here, we suggested that some functional ribosomes interacted with Upf3B composed NMD complexes in RS cells, when Upf3B-NMD complex associated ribosomes encountered 5’UTR on transcripts would activate NMD by branched UPF3B-Dependent NMD Pathway[33] and perform RNA clearance since RS do not need these RNAs, such as cell cycle related genes (Fig 8).

**Fig 7.**
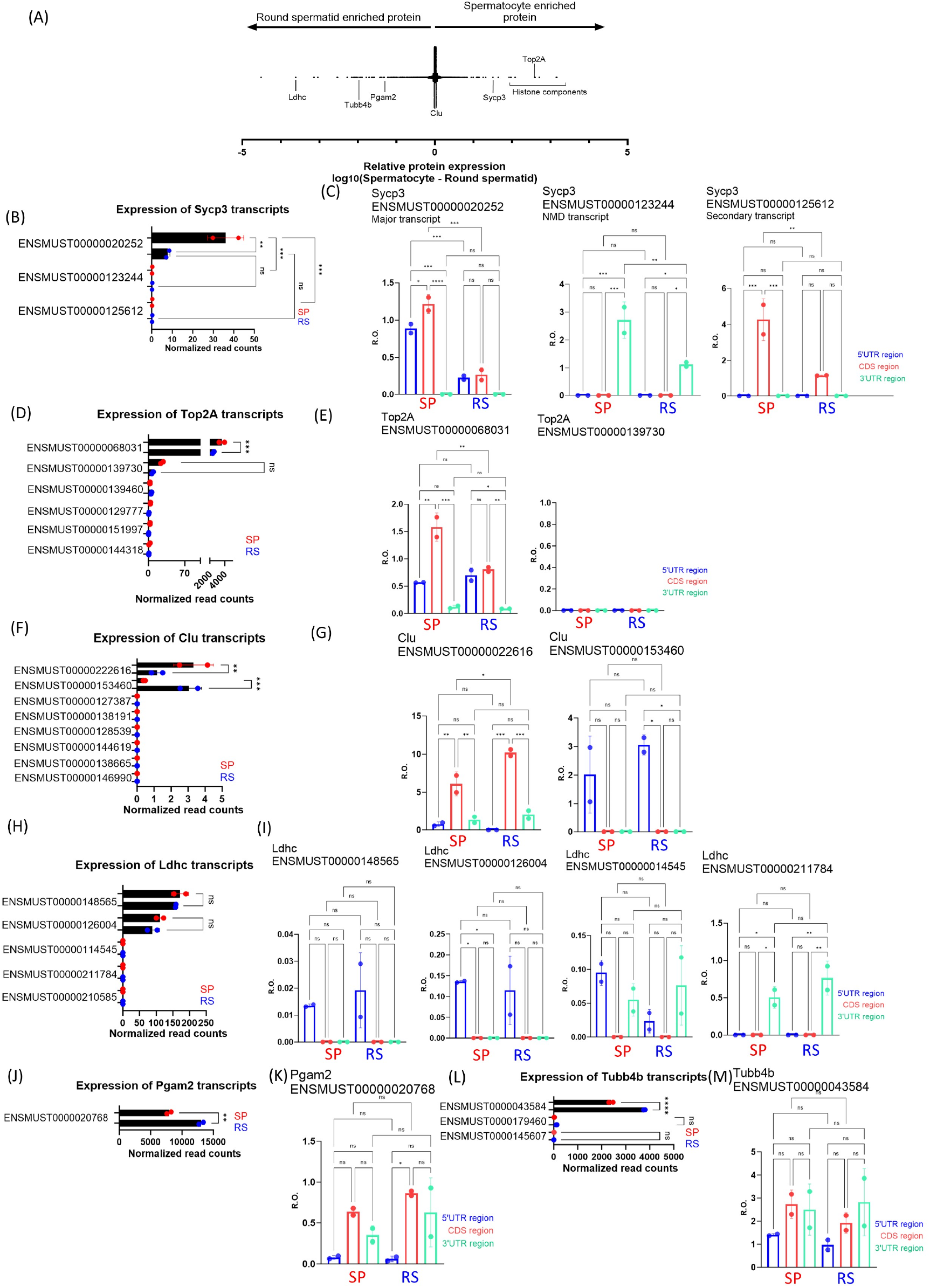
SP transcripts and proteins were downregulated when R.O. of 5’UTR in RS was dominant. (A) Comparison of proteome between spermatocytes and round spermatids. SP markers, Sycp3 and Top2a, were pinpointed. Ldhc, Pgam2, and Tubb4b were highly expressed in RS. Clu proteins were equally expressed in SP and RS. (B). RNA expression of Sycp3 different transcripts. (C). R.O from the region of 5’UTR, CDS, and 3’UTR in Sycp3 transcript. (D). RNA expression of Top2A different transcripts. (E). R.O from the region of 5’UTR, CDS, and 3’UTR in Top2A transcript. (F). RNA expression of Clu different transcripts. (G). R.O from the region of 5’UTR, CDS, and 3’UTR in Clu transcript. (H). RNA expression of Ldhc different transcripts. (I). R.O from the region of 5’UTR, CDS, and 3’UTR in Ldhc transcript. (J). RNA expression of Pgam2 different transcripts. (K). R.O from the region of 5’UTR, CDS, and 3’UTR in Pgam2 transcript. (J). RNA expression of Tubb4b different transcripts. (L). R.O from the region of 5’UTR, CDS, and 3’UTR in Tubb4b transcript.

**Fig 8.**
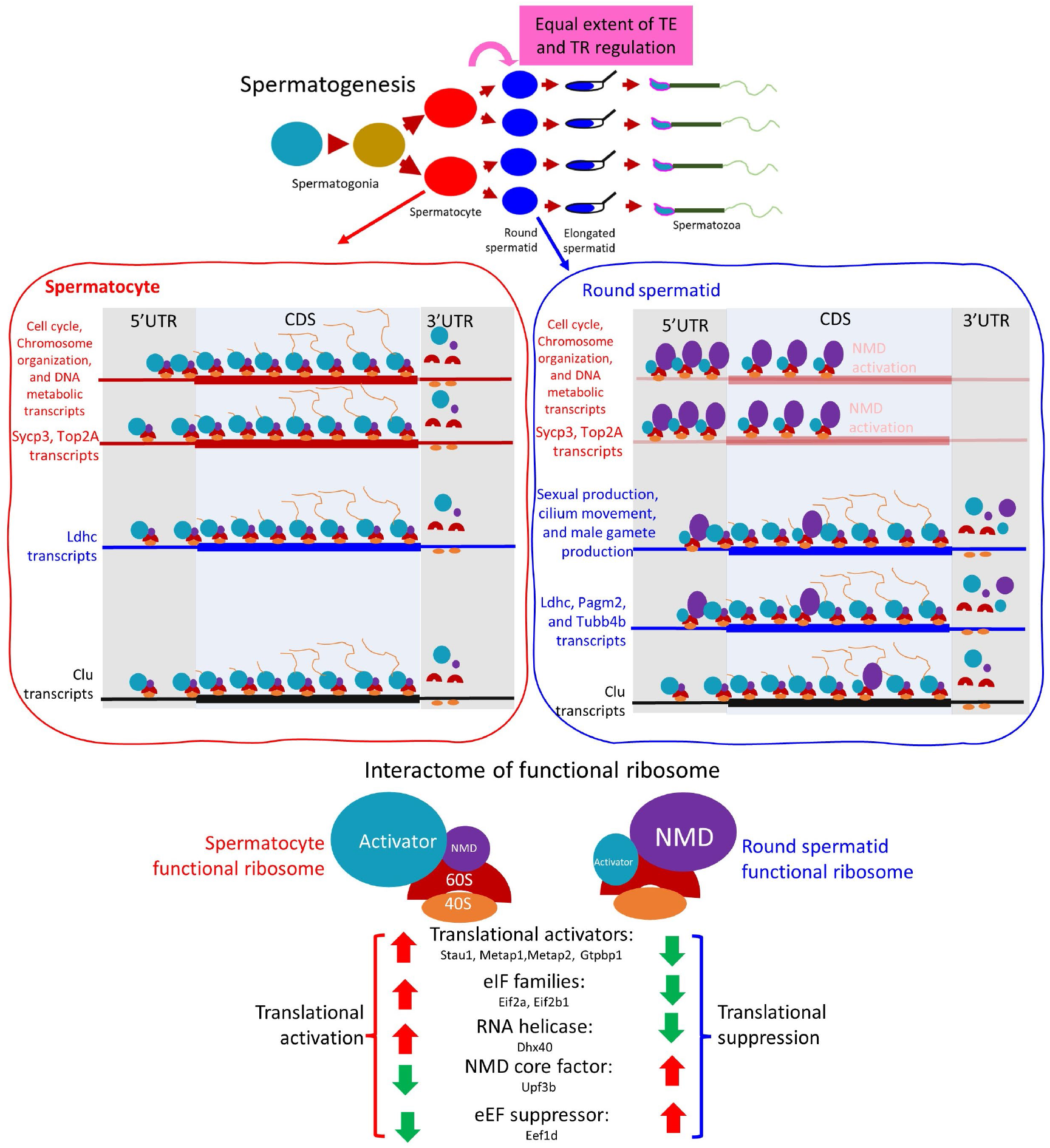
Schematic illustration of conclusion. Functional ribosomes translated cell cycle related proteins in spermatocytes which correlated with its’ mitotic/meiotic function. However, after entering round spermatid, sycp3 and cell cycle related transcripts needed to be degraded. Functional ribosomes interacted with NMD factors and accumulated in 5’UTR as in Sycp3 or both 5’UTR and 3’UTR in other cell cycle related genes. Ribosome stalling with NMD factors involvement may cause RNA degradation and explained the poor correlation of translation efficacy and proteome in round spermatids.

We claimed that transformed ribosome occupancy in SP-RS differentiation was correlated RNA clearance to avoid RS expressed SP specific proteins, such as Sycp3 and Top2a. This alteration came from the functional ribosomal interactome, NMD factors and elongated suppressor associated functional ribosomes in RS were responsible to degrade SP specific transcripts. The mature NMD complex associated ribosomes in RS may related to Upf3B, a X-linked gene, expression increasing compared to SP which is X chromosomal inactivation (XCI)[45]. The branched UPF3B-dependent NMD pathway was inactivated due to XCI in meiotic states has been known[45], but we claimed that the branched UPF3B-dependent NMD pathway was re-activated after meiosis end. There are many published data conclude that the RNA binding proteins in 3’UTR regulate the stability of RNA and facilitate RNA translation[34, 46, 47], also the length of 3’UTR and RNA modification at 3’UTR are critical factors for the microRNA regulation and translation regulation of transcripts[48, 49]. The finding here, we proposed 5’UTR may be an important regulatory element for RNA clearance by a branched NMD pathway in spermatogenesis, and answered unsolved question that RS had the poorest protein synthesis in all spermatogenic cells. There is still need in vivo experiments such as Upf3B knockout or Upf3A and Upf3B double knockout mouse models for validate the functional ribosomes of RS which occupied 5’UTR of SP specific transcripts participated the RNA degradation. However, is the branched UPF3B-dependent NMD pathway, in SP to RS transition, exon junction complex dependent or independent? The answer was unclear here. The possibility of ribosomal heterogeneity[22] in spermatogenesis was raised, based on the different interactome of functional ribosome between SP and RS. As spermatogenesis is a process depending on post-transcription regulation, we suggested spermatogenesis is a decent model for investigation of the mechanism about functional ribosome interactome and ribosomal heterogeneity.

## MATERIALS AND METHODS

### Dataset acquisition

In this paper, we collected transcriptome from RNAseq datasets, ribosome foot printing from Riboseq datasets, and proteome from quantitative mass spectrometry datasets. All these datasets were downloaded from public depositors: GEO, ArrayExpress, and ProteomeXchange.

Considering ribosome foot printing and transcriptome should be paired, we acquired these sequencing results from the following datasets, respectively. mESC leaving pluripotency (36hrs w/o LIF in medium) were extracted from E-GEOD-30839[12]. The murine brains, livers, and 4 types cells of mouse spermatogenesis were extracted from E-MTAB-7247[18]. The less committed and ground state mESC(2iL), undifferential mESC(SL), and epiblast stem cells were extracted from GSE133794[15]. The 4 cell types of neuron stem cell differential process were extracted from GSE94991[17]. Oligodendrocyte progenitor cells and oligodendrocytes were extracted from GSE182811[16]. BJ primary fibroblast cells, pre-senescent cells and senescent cells after human telomerase reverse transcriptase and tamoxifen-inducible HRAS^G12V^, and transformed cells with expressing human telomerase reverse transcriptase, p16^INK4A^-Knock-Down (KD) p53-KD and SV40 small-T were retrovirally transduced with pBabe-puro-HRAS^G12V^ were extracted from GSE45833[14]. MYC expressed U2OS and control cells were extracted from GSE66929[13]. Proteome datasets which corresponding to certain states of mESC, murine organs, and spermatogenic cells were extracted from PXD010621[50], PXD006165[51], PXD025201, PXD017284[52], supplementaryal information[53], and PXD018843[54].

### Sequencing data analysis

The majority processing from raw data to normalized read counts was operated on the public cloud platform GALAXY (usegalaxy.org)[55]. Trimmomatic package[56] was applied for reads trimming. HISAT2 [57]and featureCount [58]were used for reads alignment, assembling, and annotated by given references. Deseq2 package[59] was applied to determine the differential read counts from its normalized output. Here, we applied Deseq2 to normalize all the datasets simultaneously to avoid batch effects. Even though, there still a consideration of batch effects and unevenly biological repeats in each tissue or cell type, which were common features in datasets combination. Genome references were output from UCSC table browser, mm10 was the version we applied in each sequencing processing. Processed results of SP and RS for using in this paper was in the supplementary data 1.

### Proteome data analysis

Maxquant software was applied to process all the raw data which was downloaded from ProteomeXchange to quantitative protein expression data. Default indexes were applied to all of the processing procedures. Elongated spermatid proteome data was the exception case, which directly downloaded from its supplementaryal data. Due to proteomic data having more various divergence, most of proteome data from different publishers may not compare each other. Fortunately, the frame of SP to RS were derived from same publisher and have equal quality and depth in SP and RS dataset, which let us compare each other in this research.

### Gene Set Enrichment Analysis(GSEA)

In this paper, GSEA web tool[60, 61] was be used to describe the comparison transcriptome, R.O., and proteome between different cell types. False discover rate (FDR) was the major index to evaluate the gene enrichment in certain genesets. ssGSEA[32] were performed on GenePattern[62] platform with standard index setting.

### Mouse model and animal care

Experiments were carried out with 56-day postnatal male wildtype C57BL/6J mice. Animals were kept at room temperature with a natural night±day cycle and fed with standard laboratory chow and water ad libitum. The testis form mice were collected right after cervical dislocation. A total of 11 mice have been applied to this procedure and extracted spermatocytes and round spermatids in this paper. All animal experiments were conducted in accordance with the Guide for the Use and Care of Laboratory Animals (Animal Research: Reporting of In Vivo Experiments guidelines)[63], and all of the animal protocols have been approved by the experimental animal committee, Academia Sinica, Taiwan. The IACUC number is 21-02-1634, which is supported by MOST funding: MOST 109-2320-B-001-014-MY3.

### Interactome of functional ribosomes

Testes were collected from B6 mice when P56. SP and RS cells were isolated by STA-PUT methods[64]. The interactome of SP and RS functional ribosome were done as previous study[22]. To pull down the functional ribosomes, not the partial component, beads conjugated with Rpl36 targeted antibody (A305066AT, Bethyl) were added to cytoplasmatic fraction of SP and RS. TMT10plex™ were used to perform quantitative high-resolution MS/MS spectrum. The results were analysis by LC-MS Data Processing Software from Thermo Fisher Scientific Inc. The data were deposited to MassIVE under ProteomeXchange Consortium with accession number MSV000091932 (PXD042202 in ProteomeXchange). Processed results of SP and RS for using in this paper was in the supplementary data 2.

## Credit authorship contribution statement

S.S.L. and Y.C.K. performed the experiments and wrote the manuscript; S.S.L. conducted data analysis; Y.S.J. supported, wrote and revised the manuscript.

## ACKNOWLEDGMENTS

We thank the Common Equipment Core of IBMS and Academia Sinica of Taiwan, including the Light Microscopy Core Facility, SPF animal facility and DNA sequencing facility (AS-CFII-108-113, AS-CFII-108-103 and AS-CFII-108-115). We also thank the National Center for Genome Medicine at the Institute of Biomedical Sciences, Academia Sinica, Taiwan for technical support of the SEQUENOM MassARRAY® System.

---

**Supplement Table1.**
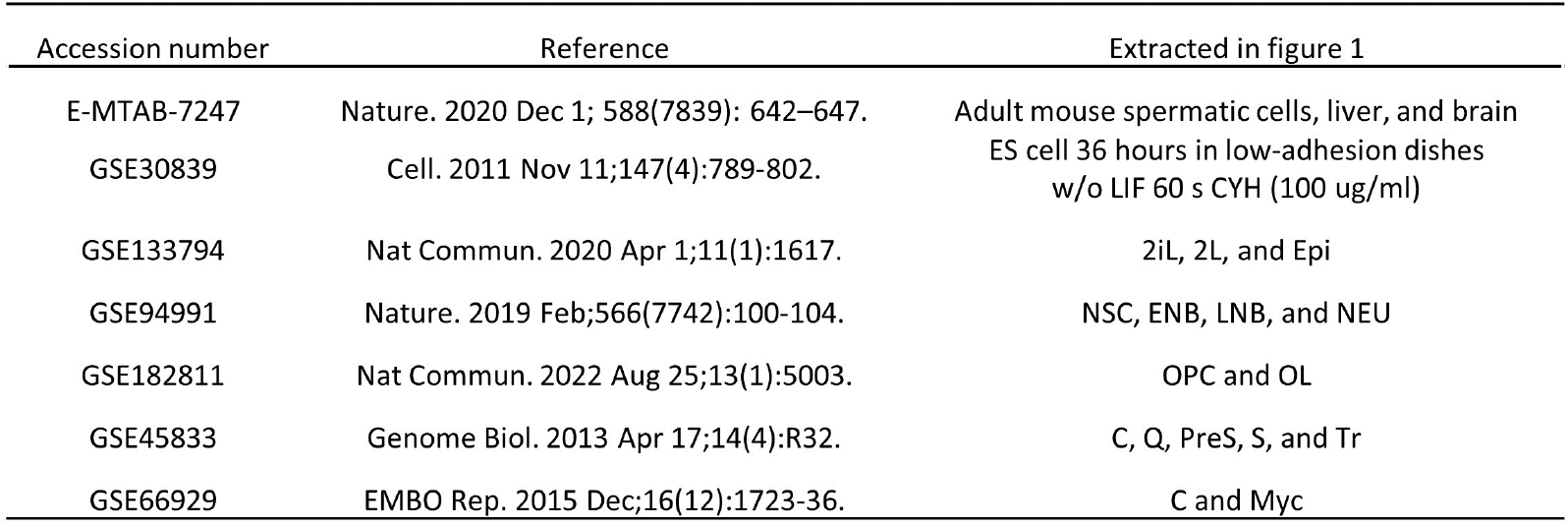
The data source of transcriptome, translatome in Figure 1.

**Supplement Table2.** The data source of transcriptome, translatome, and proteome in Figure 2 and Figure 3.

| Types | mESC<br>(w/o LIF, leaving<br>pluripotent state) | mLiver | mBrain | Spermatocyte | Round spermatid | Elongated<br>spermatid | Sperm |
| --- | --- | --- | --- | --- | --- | --- | --- |
| Transcriptome/RNAseq | GSE30839<br>, n=2 | E-MTAB-7247,<br>n=5 | E-MTAB-7247, n=3 | E-MTAB-7247, n=2 | E-MTAB-7247, n=2 | E-MTAB-7247, n=2 | E-MTAB-7247, n=2 |
| Translatome/Ribosome<br>footprinting | GSE30839<br>, n=4 | E-MTAB-7247,<br>n=5 | E-MTAB-7247, n=3 | E-MTAB-7247, n=2 | E-MTAB-7247, n=2 | E-MTAB-7247, n=2 | E-MTAB-7247, n=2 |
| Proteome/ Mass spectrum<br>result | PXD010621, n=4 | PXD006165,<br>n=10 | PXD025201, n=8 | PXD017284, n=1 | PXD017284, n=1 | PMID: 31994209,<br>suppl. Info., n=1 | PXD018843, n=1 |

**Supplement Figure 1.**
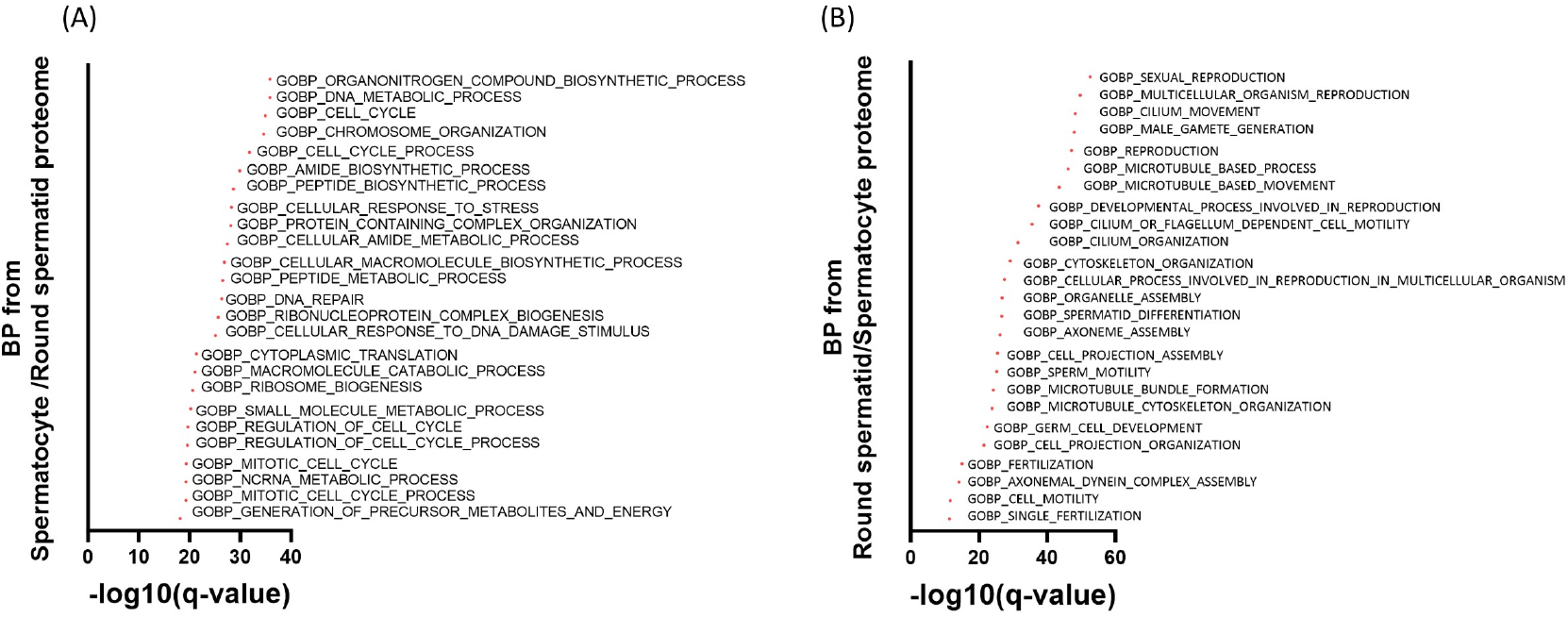
GO term represented differential expressed proteins of SP and RS which reflect the diverged biological function in these two cell types. (A)(B) Dominant Biological Process GO terms from proteome comparison between spermatocytes and round spermatids.

**Supplement Figure 2.**
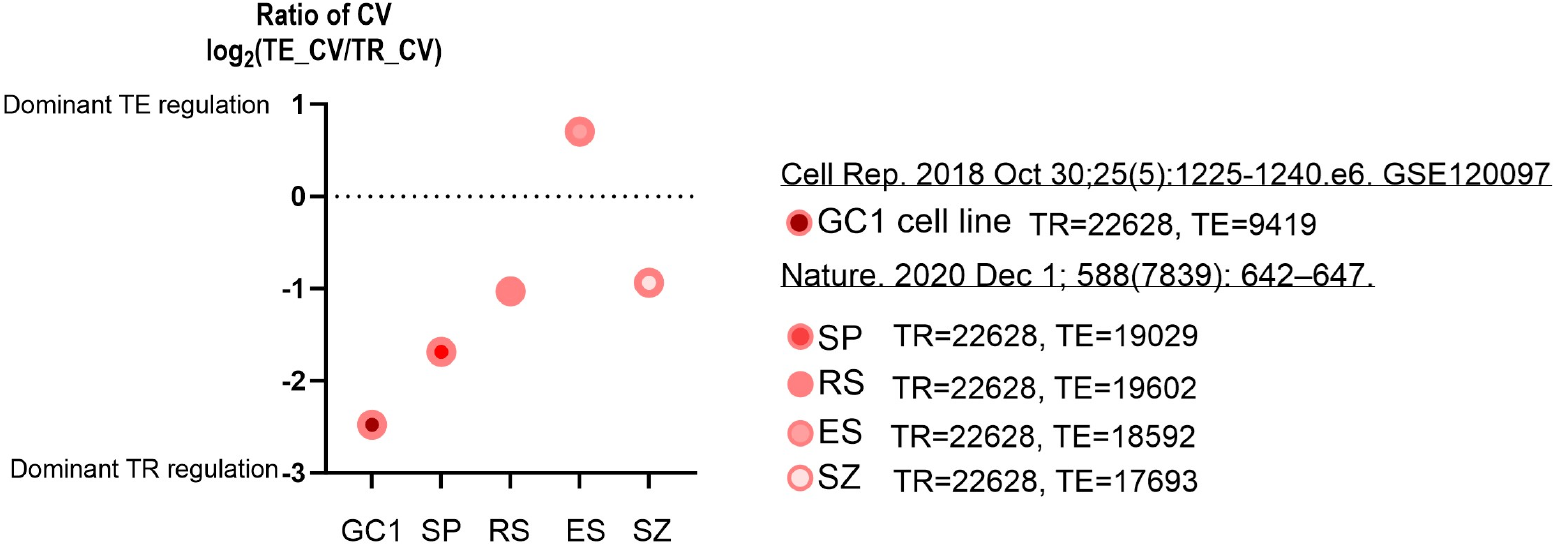
Gradually decreasing transcription regulation and increasing translation regulation through spermatogenesis. GC1: SV40 immortalized spermatogonia cell line

**Supplement Figure 3.**
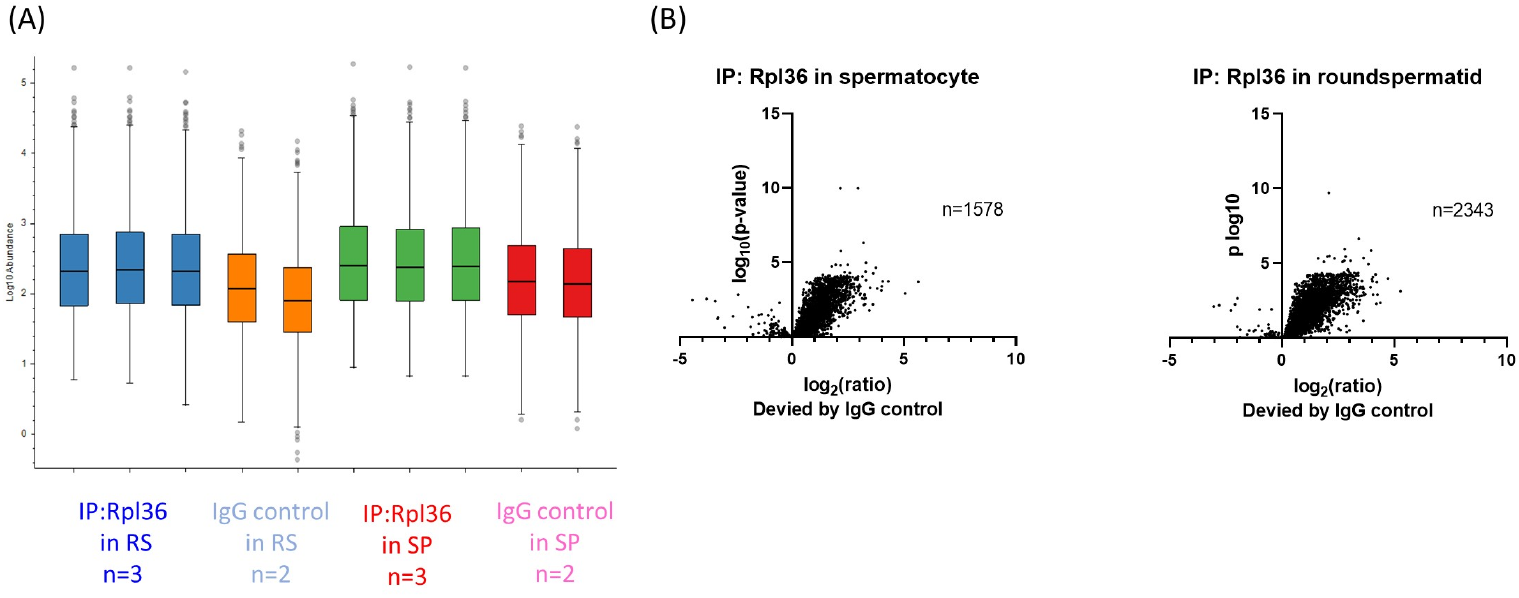
Detailed information of ribosome associated proteins (RAPs) of functional ribosomes from IP-MS. (A)Box plot of normalized input peptides. (B)Negative control (IgG) were compared to reveal actual RAPs identification.

**Supplement Figure 4.**
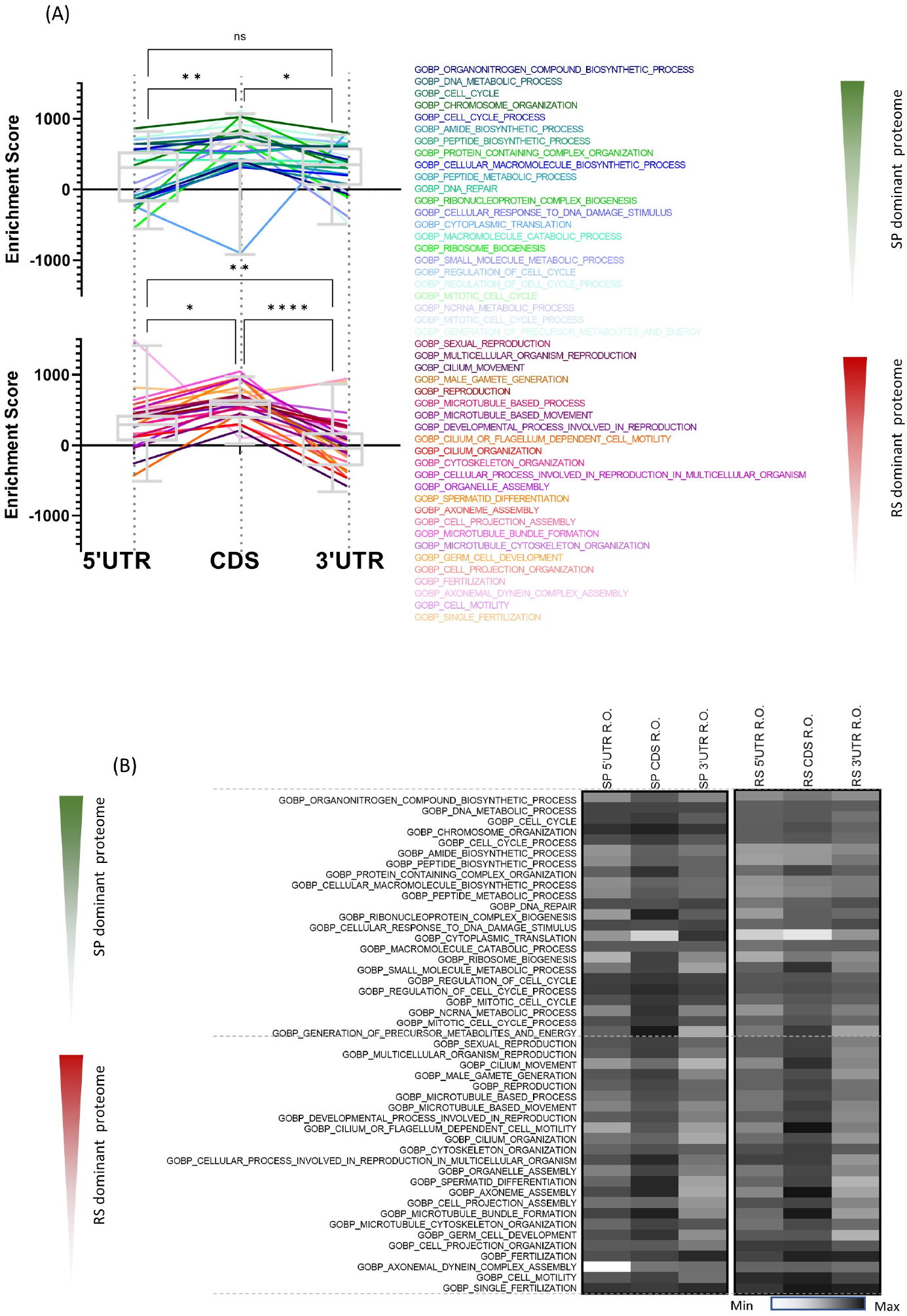
Extensional figures of Figure 5. (A) The SP proteome and RS proteome enrichment analysis of ribosome occupancy in different regions from SP.(B) Heatmap was applied to represent enrichment score of R.O. value in different regions from SP or RS.

